# Single neuron activity predicts behavioral performance of individual animals during memory retention

**DOI:** 10.1101/2020.12.30.424797

**Authors:** Martin Fritz Strube-Bloss, Tiziano D’Albis, Randolf Menzel, Martin Paul Nawrot

## Abstract

In 1972 Rescorla and Wagner formulated their model of classical Pavlovian conditioning postulating that the associative strength of a stimulus is expressed directly in the behavior it elicits^1^. Many biologists and psychologists were inspired by this model, and numerous experiments thereafter were interpreted assuming that the magnitude of the conditioned response (CR) reflects an associative effect at the physiological level. However, a correlation between neural activity and the expression of the CR in individual animals has not yet been reported. Here we show that, following differential odor conditioning, the change in activity of single mushroom body output neurons (MBON) of the honeybee predicts the behavioral performance of the individual during memory retention. The encoding of the stimulus-reward association at the mushroom body output occurs about 600 ms before the initiation of the CR. We conclude that the MB provides a stable representation of the stimulus-reward associative strength, and that this representation is required for behavioral decision-making during memory retention.

---

In classical conditioning experiments, learning performances of individual animals are often not adequately described at the group level. This applies to both, *mammals*^2^ and *insects*^3–5^. We propose that the expression of a behavioral conditioned response in the individual can be explained by the learning-induced changes in the activity of single neurons. We tested this hypothesis by recording single-unit activity in the mushroom body (MB) of honeybees while animals underwent a classical learning paradigm.

The MBs are paired structures of the insect brain that mediate high-order brain functions and memory formation^6–8^. In honeybees, each MB consists of ~170.000 Kenyon cells (KC) receiving multi-modal sensory information at the input. At this stage, sensory information converges with the reward pathway, which is facilitated by the ventral unpaired median neuron one of the maxillary neuromere (VUMmx1)^9^. The neurotransmitter octopamine mediates the reward^10,11^. Incoming olfactory information is represented by KCs in a temporally and spatially sparse code^12–14^. KC axons form synaptic connections with a relatively small population of ~400 MBONs, each of which receives converging input from several KCs^15^. MBONs typically show broad odor-response profiles and increase their response to reward-associated odors after training^16–18^.

We performed multiple single-unit recordings from MBONs in the ventral aspect of the MB vertical lobe during differential odor conditioning (Fig. 1b). Prior to conditioning (PRE), two odors were presented to the animal. During conditioning (COND), one odor (CS+) was paired with a delayed sucrose reward (US), whereas the second odor (CS-) was presented unrewarded. Three hours after conditioning, in the memory-retention phase (MEM), both odors were presented unrewarded to test whether the animals expressed the conditioned response (CR) to the CS+. The three hours delay between the COND and the MEM phases was chosen to allow the formation of a middle-term memory. In two datasets^16,17^ we co-analyzed neural and behavioral responses in all three experimental phases. To this end we computed the single-unit neural response to each single trial of odor stimulation as the baseline-corrected firing rate in the time window between 200 and 550 ms after stimulus onset. To monitor the animals’ behavior, we recorded the electrical activity of the M17 muscle, which drives the proboscis’ extension. In each single trial we detected the expression of the CR and computed the timing of its onset based on the power of the M17 signal (see Methods).

**Figure 1:**
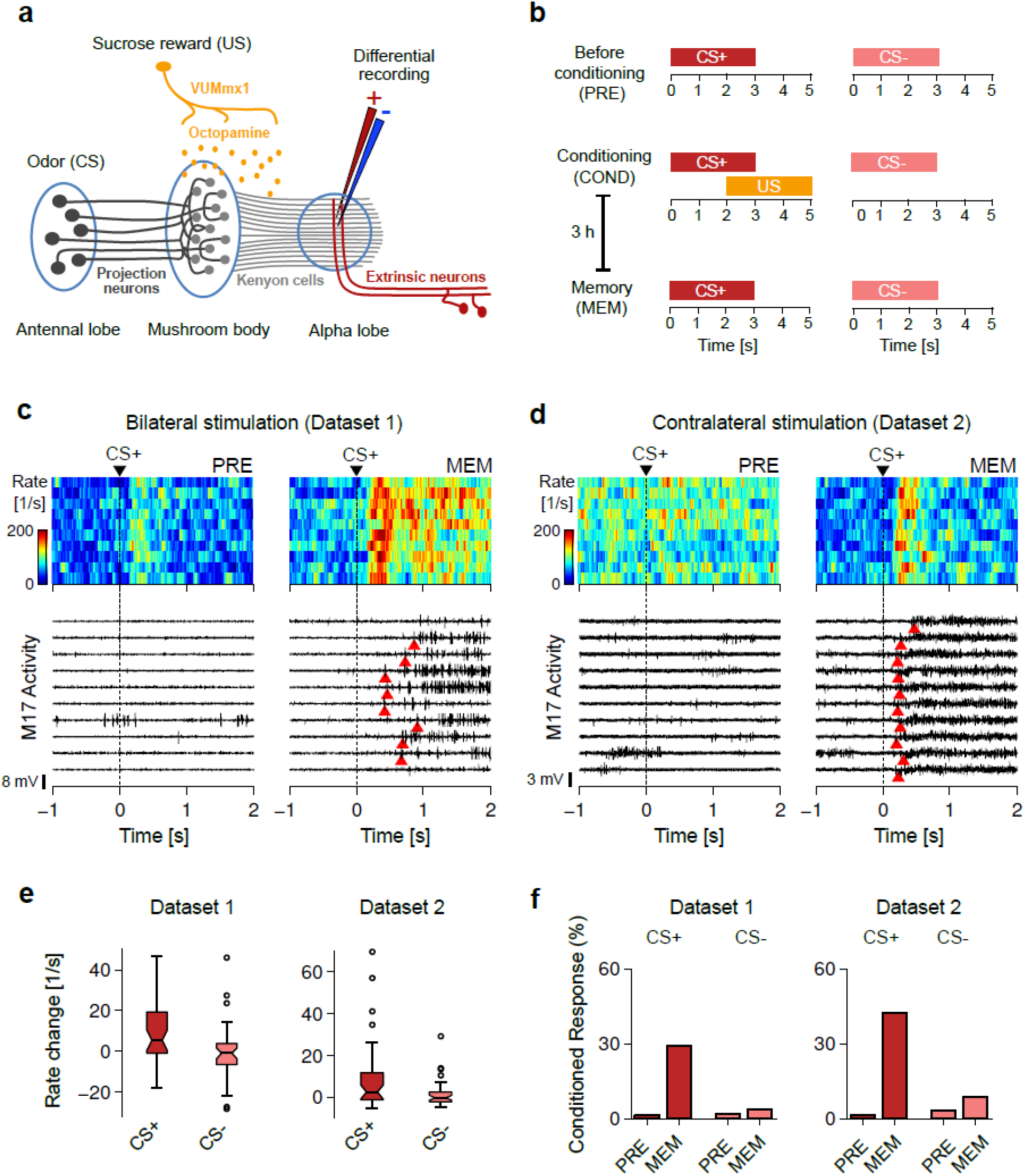
Odor-reward representation at the mushroom-body output following classical conditioning. **a**, Convergence of sensory and reward pathways at the mushroom body (MB). Projection neurons provide olfactory information to the MB and target the Kenyon cells (KC). KCs project to the ventral aspect of the alpha lobe and form synapses with mushroom body output neurons (MBONs). Reward information is conveyed by the VUMmx1 neuron and is mediated by octopamine. **b**, Experimental protocol. Two odors (CS+ and CS-) are presented 10 times in pseudo-random order before (PRE), during (COND), and three hours after conditioning (MEM). During conditioning (COND) CS+ is paired with a sucrose reward (unconditioned stimulus, US), whereas CS-is not rewarded. **c,** Examples of neural and muscular activities recorded from one animal (bee #69) subject to bilateral odor stimulation (dataset 1). The firing rate of one MBON (top) and the activity of the M17 muscle (bottom) are shown for 10 repetitions of CS+ stimulation, before (PRE, left panels) and after conditioning (MEM, right panels). Dashed lines indicate the onsets of odor stimulation. Red triangles indicate the onsets of the detected conditioned responses (CR, see online Methods for detection criteria). **d**, Same as in c but for an animal subject to contralateral odor stimulation (bee #45, dataset 2). **e**, Distributions of the trialaveraged neural responses to CS+ (red) and CS- (pink) for all the recorded units (dataset 1: N=38, dataset 2: N=29), before conditioning (PRE), and after conditioning (MEM). *: Wilcoxon rank sum test, p<0.05. **f**, Fraction of trials in which stimulation with CS+ (red) and CS- (pink) evoked a conditioned response. Trials are pooled across animals.

In many animals, differential conditioning led to a novel or increased response of MBONs in response to CS+, and to the expression of a CR in the MEM phase (Fig. 1c and d). In both datasets, neural responses to CS+ were significantly larger in the MEM phase as compared to the PRE phase (Wilcoxon rank sum test: p<0.05, Fig. 1e), and the same effect was not observed for CS- (Wilcoxon rank sum test: p>0.4). Similarly, the number of CRs increased considerably from the PRE to the MEM phase following CS+ stimulation, but remained unchanged following CS-stimulation (Fig. 1f).

To quantify the relationship between CS+ response and CR we analyzed the dynamics of neural and behavioral responses on a trial-by-trial basis. Figure 2a shows examples of trial-resolved neural responses to CS+ (solid lines) and CS-(dashed lines) for all recorded units in four animals (one animal per row), and the detected CRs in the corresponding trials (black dots: CS+ trials, gray dots: CS-trials). Most of the animals (dataset 1: 10/17, dataset 2: 9/14) developed a CR to CS+ already in the conditioning phase (COND), despite neural responses to CS+ did not change significantly between the PRE and COND phases (Wilcoxon rank sum test, dataset 1: p>0.3, dataset 2: p>0.9, see for example bees #69, #83 and #67). This effect was also evident at the population level, when trial-resolved neural and behavioral responses were averaged across animals (Fig. 2b, top): following CS+ stimulation, the behavioral learning curve (gray line) gradually increased during training (COND), but the average neural response (red line) increased abruptly only with the first trial of the MEM phase. This suggests that activity changes of MBONs reflect consolidation-dependent memory rather than short-term memory, as in the latter case enhanced responses to CS+ should have been observed starting with the COND phase. As a control, we further verified that both the behavioral learning curve and the average neural responses of MBONs remained nearly flat across the three phases of CS-stimulation (Fig 2b, bottom).

**Figure 2:**
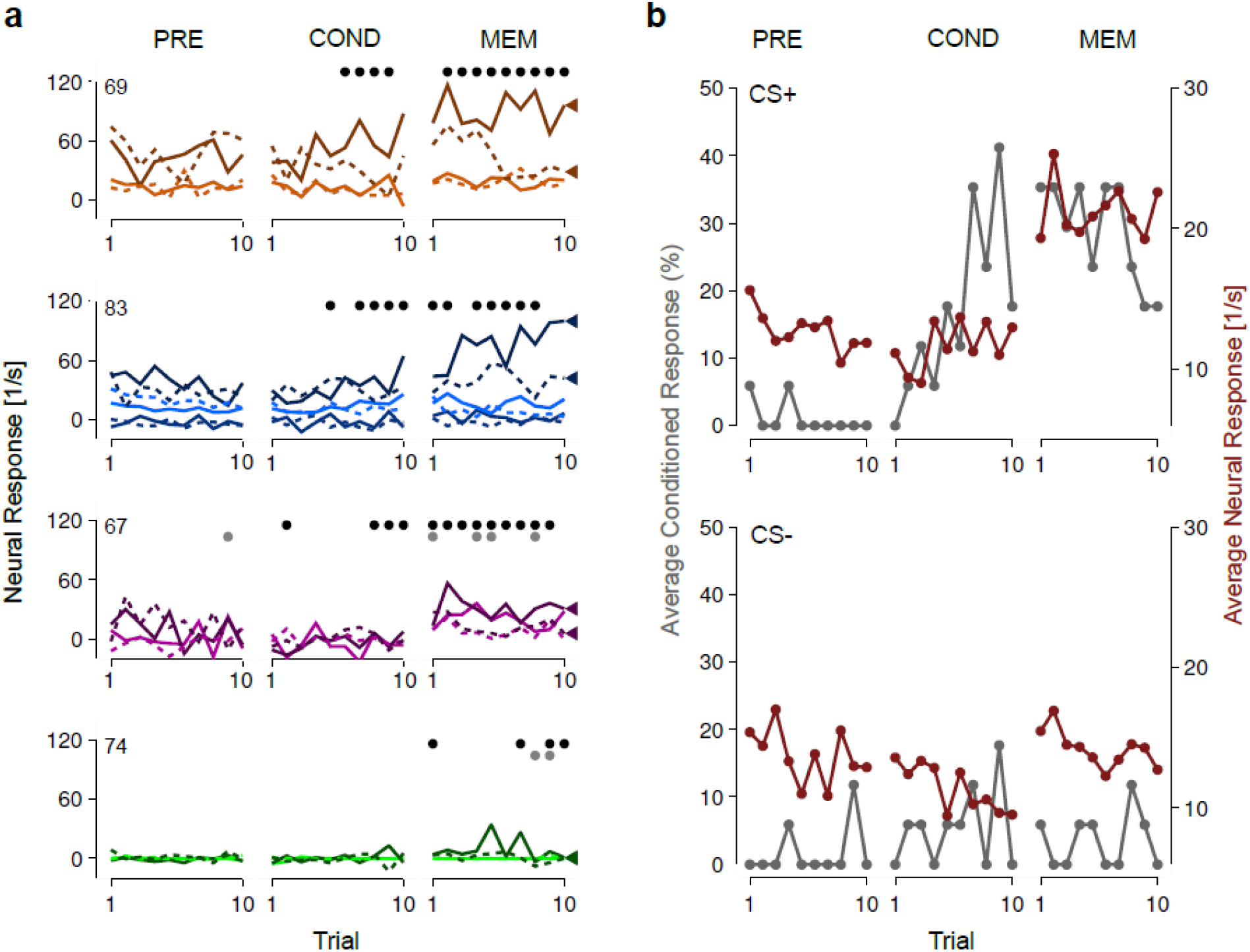
Trial-resolved dynamics of neural and behavioral responses. **a**, Neural responses (lines) of all the recorded units from four example animals in dataset 1 (one animal per row, animal IDs: 69, 83, 67, 74), before (PRE), during (ACQ) and after conditioning (MEM). Each color corresponds to a single unit, solid (dashed) lines indicate responses to CS+ (CS-). For each animal, the triangles on the right-hand side indicate the unit where the difference between CS+ and CS-neural responses in the MEM phase increased maximally as compared to the PRE phase. Black (gray) dots indicate behavioral CRs to CS+ (CS-). **b**, Population dynamics of neural and behavioral responses (dataset 1). Top: Fraction of animals that expressed a CR in a trial of CS+ stimulation (gray), and corresponding neural responses averaged across all units of all animals (N=38). Bottom: same as in the top panel but for trials of CS-stimulation.

Interestingly, we observed that good learners, i.e., animals that learned to better discriminate between CS+ and CS-behaviorally, also showed larger differences between CS+ and CS-neural responses during the MEM phase (compare between examples in Fig. 2a). We then asked whether the learning-induced change of neural response in an individual is predictive of its behavioral performance during memory retention? We quantified the changes in neural and behavioral responses by computing, for each animal, the conditioned-plasticity (CP) score and the conditioned-response (CR) score. The CP score measures the difference between CS+ and CS-neural responses in the MEM phase as compared to the PRE phase. Similarly, the CR score measures the difference between CS+ and CS-behavioral responses in the MEM phase as compared to the PRE phase (see Methods for details). Strikingly, the two scores correlated strongly and significantly across individuals in both datasets (pairwise linear correlation coefficient, dataset 1: r=0.93, p<0.001, Fig. 3a; dataset 2: r=0.59, p<0.05). This establishes for the first time a direct link between the learning-induced activity changes of MBONs and the learning-induced behavioral expression of a CR in the single animal.

**Figure 3:**
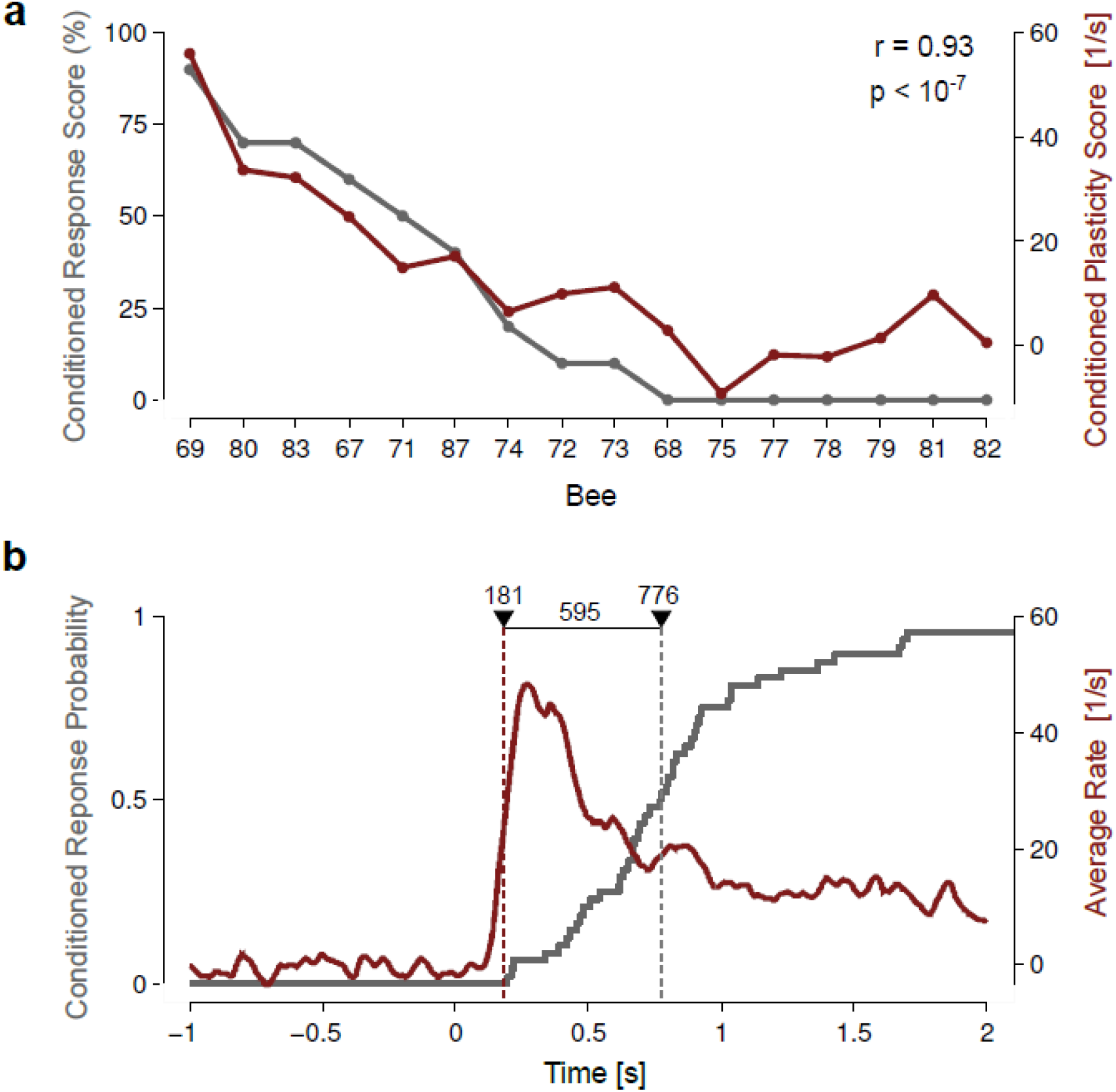
Change in neural activity correlates with behavioral performance across individuals. **a**, Correlation between changes in neural activity (conditioned-plasticity score, red line) and behavioral performances (conditioned-response score, gray line) for all the animals in dataset 1. Data-points are ordered according to the conditioned-response score. A similar correlation is found for dataset 2 (r=0.59, p<0.05). **b**, Temporal dynamics of neural and behavioral responses within a trial of odor stimulation (dataset 1). Red lines: average activity of all MBONs that responded to CS+ during the MEM phase (red solid line), and median onset of the same responses (181 ms, red dashed line). Gray lines: cumulative distribution of the onset times of all detected CRs to CS+ during the MEM phase, and median onset of the same responses (776 ms, gray dashed line). The difference between the onsets of behavioral and neural responses is 595 ms. Time zero corresponds the odor-stimulation onset.

To test whether MBON activity represents the neural memory trace that may causally underly the CR expression we quantified the temporal dynamics of neural and behavioral responses within a trial of odor stimulation (Fig. 3b). The average neural response to CS+ presentation in the MEM phase peaked about 200 ms after stimulus onset. The median onset time across individual CS+ responses was 181 ms whereas behavioral responses were delayed by about 600 ms. This relatively long delay suggests that the activity of MBONs does not trigger behavior directly, but rather supports decision making in downstream structures by contributing information about stimulus value^19^.

Our data are consistent with the view that the MB creates a high-dimensional multi-modal sensory space at the input (encoded by the activity of thousands of KCs) into a low dimensional value-based space at the output (encoded by a few hundreds of MBONs)^17^ categorizing multimodal information^20^. Here, we provide evidence that this value-based information is of direct behavioral relevance. Indeed, neural activity changes at the MB output accurately predicted behavioral performances in single animals three hours after conditioning. The fact that no significant activity changes were detected during conditioning where CRs were already expressed suggests that MBONs are necessary for the retention of a memory but not for its initial acquisition and therefore biases memory-based action selection as suggested for drosophila^21–23^. Our results contribute to the view that, in insects, separate neural pathways mediate memory acquisition and memory recall^6,24^, and that consolidation processes could transfer behaviourally-relevant information across neural structures^17^.

## ACKNOWLEDGEMENTS

This work was funded by the German Research Foundation in parts through the DFG grant (STR 1334/3-1) to MSB and through the Research Unit *Structure, Plasticity and Behavioral Function of the Drosophila mushroom body* (DFG-FOR 2705, grant no. 403329959). Experimental data sets had been funded by Bundesministerium für Bildung und Forschung (BMBF) through grant 01GQ0941 to the Bernstein Focus Learning and Memory in Decision Making.

## METHODS

Details about animal preparation, differential odor conditioning, data acquisition and data pre-processing as well as visualization of the recording electrode position were described in detail in previous publications^14,15,21,22^.

### Electrophysiology

The activity of MBONs was recorded extracellularly from the ventral aspect of the vertical-lobe at a depth between 100 and 250 μm. Axo-dendrites of MBONs are about ten times larger compared to KCs axons^13^ and thus MBON spikes could be reliably discriminated from KC spikes^14^. The activity of the M17^20^ muscle, which innervates the bee’s proboscis, was simultaneously recorded.

### Analysis of neural data

Instantaneous firing rates were estimated by a convolution of the binary spike trains with a causal exponential kernel (σ=50 ms). Single-trial neural responses were computed by averaging the instantaneous firing rates between 200 and 550 ms after stimulus onset, and by subtracting the baseline rate for the trial (average rate in the 2 s preceding odor onset). The conditioned-plasticity (CP) score was the maximum unit-plasticity (UP) score across all units recorded from a given animal. The UP score was defined as follows:

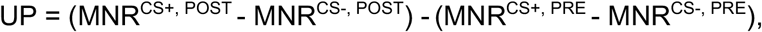

where MNR^X,Y^ is the mean neural response to odor X in experimental phase Y. The onset of a neural response (Fig. 3b) was the first time point where the baseline-corrected firing rate was two times larger than the baseline SD (SD of the firing rates in the 2 s preceding odor onset).

### Analysis of M17 data

The instantaneous power of the M17 activity was estimated by a convolution of the M17 raw signal with a causal rectangular kernel (width = 2 s). A conditioned response (CR) was detected when the instantaneous signal’s power in the M17 channel crossed a given threshold. Two thresholds were estimated for each animal: a threshold T_PRE_ for detecting responses during the PRE and ACQ phases, and a threshold T_MEM_ for detecting responses during the MEM phase. This was due to non-stationaries of the M17 signal after the 3 h delay between ACQ and MEM phases. The two thresholds T_PRE_ and T_MEM_ where estimated automatically from the following three distributions of the signal power: D_BASE,PRE_, D_BASE,MEM_ and D_RESP_. The two baseline distributions D_BASE, PRE_ and D_BASE, MEM_ were the distributions of the signal power in a 5 s time window before stimulus onset, across all trials in the PRE and MEM phases respectively. The response distribution D_RESP_ was the distribution of the signal power in the 3 s time window in which the sucrose reward was provided, across all rewarded ACQ trials. Distributions were estimated by Gaussian kernel convolution with reflection. The threshold T_PRE_ and T_MEM_ were the intersections between the response distribution D_RESP_ and the two baseline distributions D_BASE,PRE_ and D_BASE,MEM_ respectively. These thresholds correspond to the optimal decision boundaries that maximize the percent correct for an unbiased observer. The conditioned-response (CR) score was defined as follows:

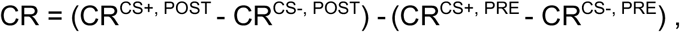

where CR^X,Y^ is the fraction of trials in experimental phase Y where odor X was presented and a conditioned response was detected.

